# Design and characterization of broadly protective influenza A(H3N2) vaccine candidates using protein language models

**DOI:** 10.64898/2026.08.26.747087

**Authors:** Victoria R. Howard, James D. Allen, Matthew H. Thomas, Giuseppe A. Sautto, Ted M. Ross, Ivelin S. Georgiev

## Abstract

Seasonal influenza A viruses cause significant global morbidity each year. Although vaccination remains the primary preventive strategy, effectiveness is often reduced by antigenic drift. This challenge is particularly pronounced for influenza A(H3N2), which has required eight vaccine updates over the past decade. Here, we present a computational framework to engineer broadly reactive influenza A(H3N2) vaccines, using protein language models to generate novel hemagglutinin (HA) sequences and a machine learning model to predict antigenic distance from circulating strains. In a proof-of-concept study, seven HA candidates designed using sequence data from 2013–2018 were evaluated in mice against contemporary and subsequently circulating viruses. Two candidates elicited protective levels of reactive antibodies, robust H3-specific antibody-secreting cell responses, and cross-neutralization against contemporary clades and drifted 2019-2020 strains. These findings demonstrate that an integrated generation–selection strategy can enhance vaccine coverage across current and future A(H3N2) seasons and may be applicable to other influenza subtypes.

## Introduction

Seasonal influenza viruses cause substantial global morbidity and mortality, resulting in ∼290,000 to 650,000 deaths annually worldwide^1^. Although vaccination remains the most effective preventative strategy, vaccine effectiveness varies considerably between seasons^2^. A major contributor to this variability is antigenic drift, whereby mutations accumulate in the hemagglutinin (HA) and neuraminidase (NA) glycoproteins, enabling escape from existing immunity and increasing the risk of vaccine mismatch^3,4^. Seasonal vaccine strains are selected approximately six months before distribution based on prior surveillance and predictions of future viral prevalence^5,6^. However, continued viral evolution between strain selection and vaccine deployment can reduce vaccine effectiveness^7,8^. The impact of this challenge was evident during the 2025–2026 influenza season, when an estimated 340,000 hospitalizations and 21,000 deaths occurred in the United States, largely driven by the emergence of an antigenically distinct H3N2 lineage after vaccine strain selection ^9,10^. These limitations highlight the need for new vaccine design strategies that better anticipate influenza virus evolution.

To address this challenge, several computational approaches have been developed to broaden vaccine derived immunity. The Computationally Optimized Broadly Reactive Antigen (COBRA) methodology generates layered consensus antigens from genetically related HA sequences, incorporating epitopes from multiple circulating strains to improve cross-reactivity within influenza subtypes^11–15^. Other approaches including Mosiac and Epigraph, use algorithm driven optimization to maximize epitope coverage across viral populations^16–19^. While these approaches effectively capture dominant patterns of observed sequence diversity and can improve protection against future variants, they are limited to the variation present in available datasets and may not capture higher-order dependencies between residues that shape antigenicity and function.

Recent advancements in machine learning (ML) demonstrate that protein language models (pLMs), trained on millions of diverse protein sequences, learn fundamental biological information by developing a statistical understanding of evolutionary patterns^20–22^. These models can be further fine-tuned to generate sequences within specific protein families that are both diverse and consistent with known functional and evolutionary properties^20^. This capability makes pLMs well-suited for exploring regions of HA sequence space, and generating candidate antigens that incorporate evolutionary patterns across influenza seasons to achieve broader effectiveness.

A complementary challenge is identifying candidate antigens with the greatest potential to provide broad protection against circulating and future viruses. Numerous computational approaches have been developed to predict influenza antigenicity from sequence data, but most rely on manually engineered features or predefined assumptions regarding antigenically important residues^23,24^. Recent pLMs trained on evolutionary trajectories, implicitly capture the structural constraints of the HA protein and infer antigenic properties directly from the protein sequence without explicit feature engineering^25,26^. Building on these advances, we used pLM-derived sequence representations to train an antigenicity prediction model that prioritizes HA candidates for experimental validation, establishing a unified pLM-based framework for both generation and selection.

To evaluate this approach, the antigenically drifted 2018–2019 H3N2 season, during which vaccine efficacy was ∼29%, was used as a historical test case^27^. A generative pLM was fine-tuned on HA sequences from viruses circulating between 2013 and 2018 to generate novel HA antigens, which were subsequently prioritized using the accompanying predictive model to downselect candidates. This process resulted in seven lead candidates that were evaluated *in vivo* for humoral, cellular, and protective immune responses. Collectively, these studies demonstrate that pLM-guided antigen design can generate vaccine candidates that elicit broad and protective immune responses against both contemporary and antigenically drifted H3N2 viruses.

## Results

### Fine-tuned pLM captures the evolutionary landscape of influenza A(H3N2) HA sequences

The generative pLM ProGen2^20^ was fine tuned on a dataset of human A(H3N2) HA sequences circulating between 2013 and 2018^28^ **(Fig. 1A)**. Sequences were restricted to the HA1 subunit, which contains the primary antigenic sites, and clustered to remove near-duplicates, yielding 701 representative sequences. To understand how exposure to successive influenza seasons influences model learning, a time-resolved validation framework was implemented in which the training set was cumulatively expanded and performance was evaluated on the subsequent season **(Fig. 1B)**. Model performance was quantified using perplexity, which measures how well a model predicts sequence data, with lower values indicating better predictive accuracy. Fine-tuning consistently reduced perplexity across all validation folds relative to the original model, demonstrating stable performance as successive seasons were incorporated into training **(Fig. 1C)**. The final model, trained on all sequences from 2013-2018, was subsequently evaluated using a held-out test set of strains isolated between 2019 and 2023. The fine-tuned model outperformed the baseline across all future seasons, demonstrating robust generalization to viruses that emerged after the training period. However, perplexity increased for more distant seasons, consistent with increasing evolutionary divergence and antigenic drift **(Fig. 1D)**.

**Fig. 1.**
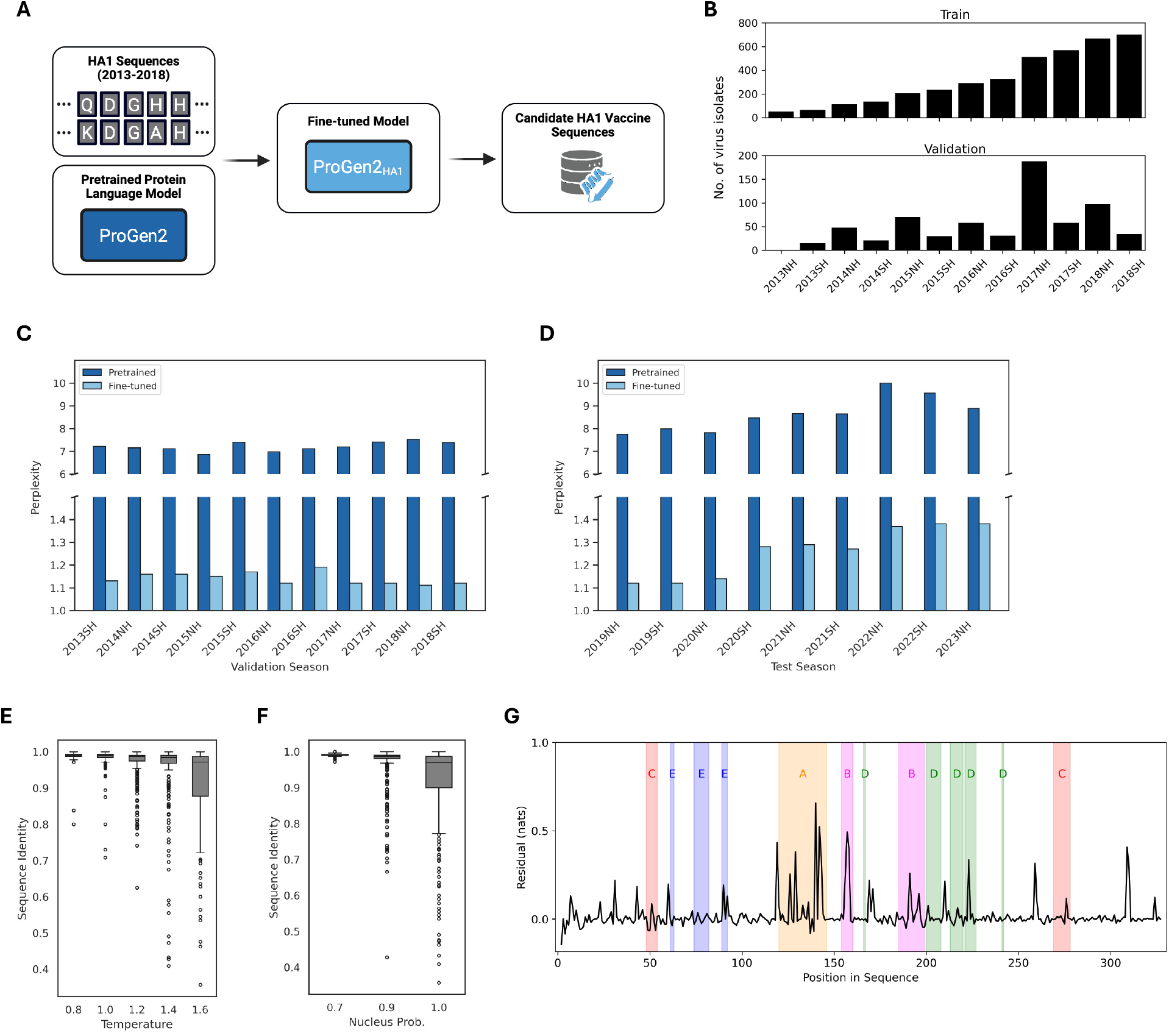
Fine-tuning ProGen2 to generate influenza A(H3N2) HA1 sequences. (A) A dataset of sequences from 2013-2018 was curated to fine-tune ProGen2 for generation of vaccine candidates. (B) Seasonal division of data into training and validation datasets respectively for training and evaluation of the fine-tuned model in a time-series cross validation scheme. (C) Perplexity (unitless) for seasons held out from each time-series fold, for both the pretrained model and the fine-tuned model. (D) Perplexity (unitless) for seasons past the training set, for both the pretrained model and the fine-tuned model. (E) Designs generated under each temperature value are plotted against their maximum sequence identity to the training set. (F) Designs generated under each top *p* value (nucleus probability) are plotted against their maximum sequence identity to the training set. (G) Residual predictive entropy (nats) at each sequence position, calculated by subtracting a centered rolling median (window = 11 residues) from the average predictive entropy across 356 filtered candidate sequences. Shaded regions indicate known antigenic sites.

### Generated HA1 sequences are diverse and evolutionarily plausible

The final model generated 1500 candidate HA sequences across 15 combinations of sampling temperatures and nucleus sampling probabilities. As expected, increasing sampling parameters increased sequence novelty, enabling exploration of diverse regions of HA sequence space **(Fig. 1E-1F)**. Initial filtering removed redundant sequences, low-quality designs, and candidates lacking sequence novelty relative to the training set, resulting in 356 remaining candidates. Localized peaks in predictive entropy coincided with several known antigenic regions, particularly sites A and B, suggesting that the model captures aspects of the evolutionary flexibility associated with these regions **(Fig. 1G)**.

### Antigenic model predicts strain relationships

To further filter candidate designs, a predictive model of antigenic distance was developed using a curated hemagglutination inhibition (HAI) dataset^29^. The dataset includes 3,698 paired A(H3N2) strains from 1968-2010 with experimentally measured antigenic distances quantified as Archetti-Horsfall titers (AHT), a measure that has been shown to correlate with vaccine effectiveness^30^. The model used HA1 amino acid sequences of virus–reference strain pairs as inputs and predicted their antigenic distance, enabling direct comparison to experimental readouts. Sequence features were extracted by the widely used pLM ESM-2^31,32^, which was fine-tuned to the dataset of HA1 sequences from 2013-2018 using parameter-efficient fine-tuning^33^. The sequence-level embeddings for both strains were then substracted and passed through a multi-layer perceptron (MLP) **(Fig. 2A)**.

**Fig. 2.**
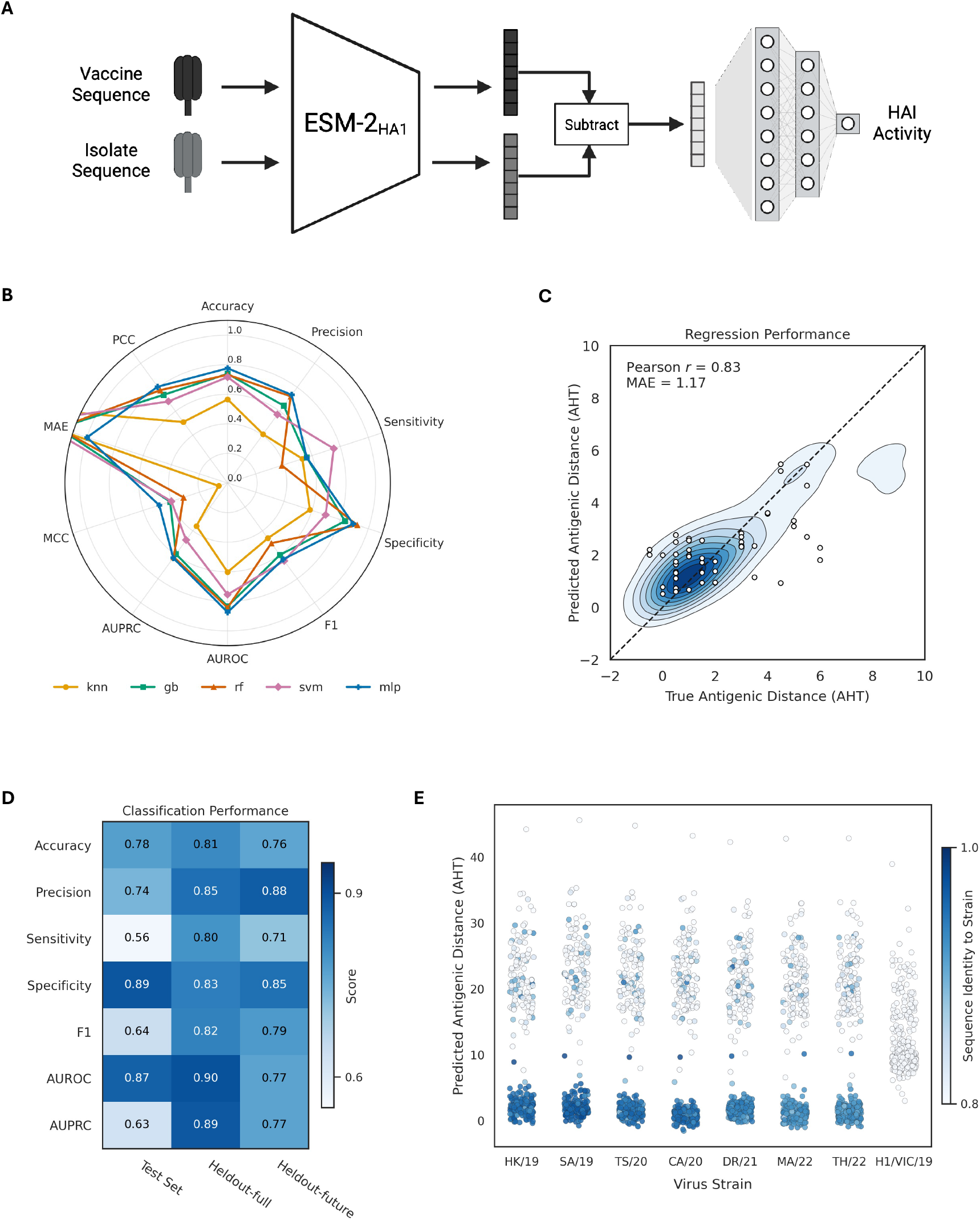
Performance of the antigenicity model for antigenic characterization of influenza A(H3N2) viruses. (A) Overview of model architecture integrating ESM-2 protein features with downstream antigenic predictions. (B) Comparison of k-nearest neighbors (KNN), gradient boosting (GB), random forest (RF), support vector machine (SVM), and multi-layer perceptron (MLP) classification and regression metrics. (C) Contour plot summarizing the performance of the MLP model on an independent holdout set of 882 AHT measurements. Antigenic pairs from the “future” (2010-11) are explicitly plotted. (D) Classification performance of MLP model on the test set, the independent holdout set, and only “future” pairs in the holdout set (2010-2011 pairs). (E) Model predictions for the filtered set of designs (n=356) against each of seven future-drifted strains, colored by the sequence identity to the strain.

The MLP was benchmarked against K-nearest neighbors (KNN), gradient boosting (GB), random forest (RF), and support vector machine (SVM) models. Hyperparameters for each model were optimized using an expanding-window time-series cross-validation scheme, in which antigenic relationships within each validation year were predicted using data from preceding seasons. The optimized models were then evaluated on a holdout set of sequence pairs from 2010. The MLP achieved the best performance across both classification and regression metrics **(Fig. 2B)**. To assess external generalizability beyond the model selection dataset, the final MLP was evaluated using an independent dataset of 882 AHT measurements spanning 1968–2011, compiled by Bedford et al.^34^. The model demonstrated strong performance on these unseen sequence pairs (MAE = 1.17, PCC = 0.83; **Fig. 2C**). To further evaluate temporal generalization, the external dataset was restricted to measurements collected after 2009. Across datasets, the model maintained strong discriminative ability (AUROC, 0.77–0.90; **Fig. 2D**) and consistently high specificity (0.83–0.89; **Fig. 2D**), supporting its use as a screening tool for prioritizing experimental candidates. However, sensitivity declined on the temporally distant subset, indicating reduced performance for comprehensive identification of all positives.

### Model-guided selection identifies diverse candidate vaccine strains with predicted broad protection

The remaining 356 candidates were evaluated against seven antigenically relevant A(H3N2) strains circulating between 2019 and 2022, together with one A(H1N1) strain as a negative control. Predicted antigenic distances (AHTs) were computed for each candidate–strain pair using the antigenicity model. Candidates predicted to be antigenically similar to a given strain generally exhibited higher sequence identity, whereas all candidates were predicted to be antigenically distant from the A(H1N1) control. Consistent with ongoing viral evolution, sequence identity also decreased for strains increasingly distant from the training period (2013–2018) **(Fig. 2E)**.

Candidates were then prioritized in a two-stage selection process. First, only candidates predicted to provide protection against all seven future A(H3N2) strains were retained, yielding 75 broadly reactive designs. Second, a diverse subset of 30 candidates was selected using Levenshtein distance within antigenic regions to maximize exploration of antigenic sequence space while minimizing redundancy.

### The candidate HA amino acid sequenced shared a common homology with the SG/16 isolate

From the 30 HA sequences, seven candidate HAs were selected based on their phylogenetic relationships to historical A(H3N2) vaccine strains and previously characterized COBRA H3 HA antigens ^14,35^. Candidate 1035 clustered most closely with clade 3c.3a viruses SW/13 and KS/17, whereas candidates 814 and 1025 aligned with the 3c.2a strain SG/16. Candidates 536 and 585 were most similar to the future drifted strains HK/19 and SA/19, while candidates 547 and 848 were selected as intermediate sequences between SW/13 and HK/14 and SG/16 and HK/19 respectively (**Fig. 3A**). Consistent with these phylogeneitic relationships, all seven candidates HAs were most similar to SG/16, possessing 6 to 16 amino acid differences, and most most divergent from the future-emerging isolate DR/21, ranging from 24 to 31 amino acid differences (**Fig. 3B**).

**Fig. 3.**
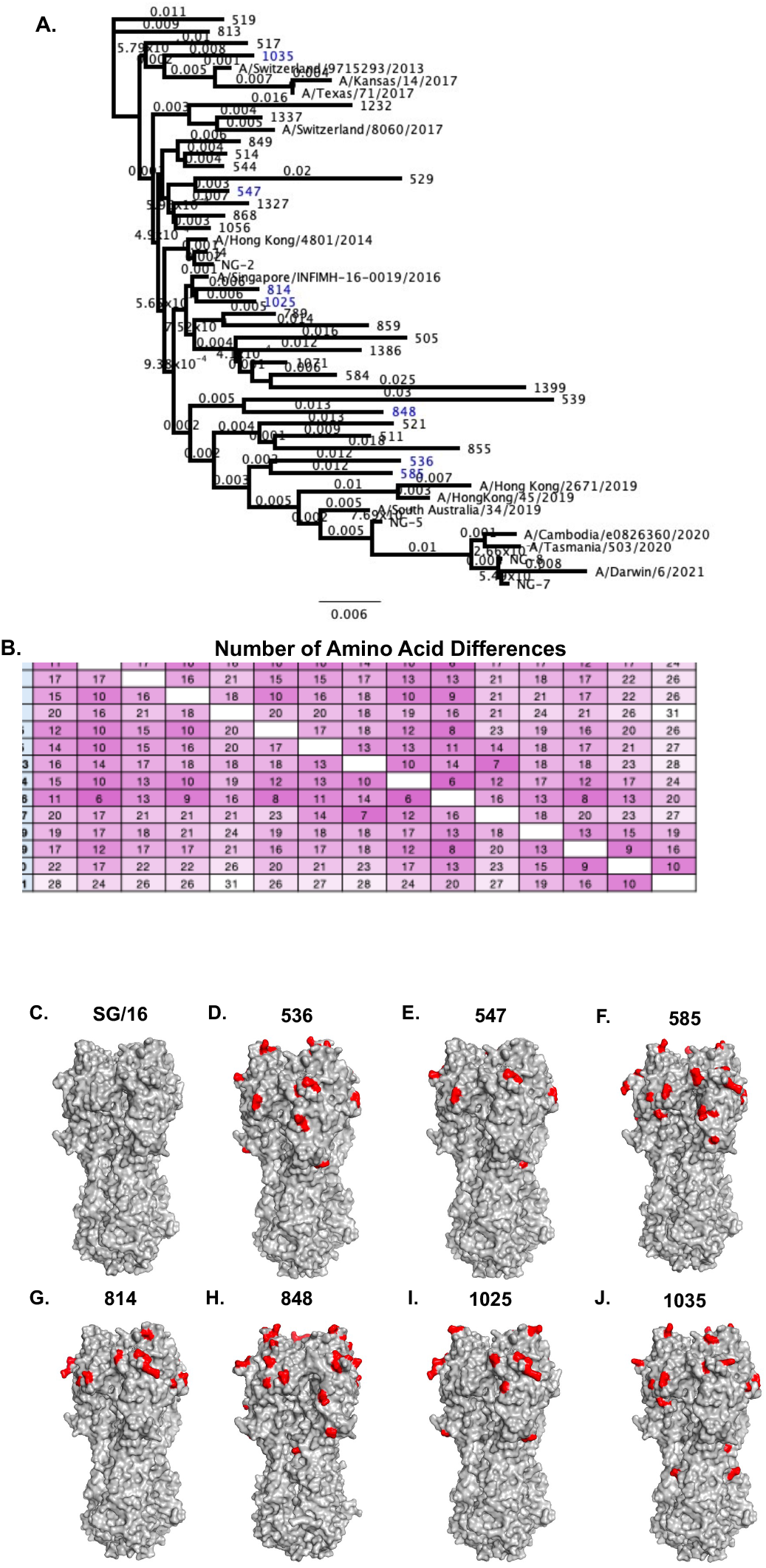
Phylogenetic and structural comparison of pLM-generated and wild-type H3 HAs. Full-length HA amino acid sequences from seven pLM-generated candidates and historical A(H3N2) vaccine strains (2013-2021) were compared using Geneious Bioinformatics software. A neighbor-joining phylogenetic tree was generated using the Jukes-Cantor distance model to assess genetic relationships among the pLM-derived and the wild-type HAs (A). Branch labels indicate substitutions per site. A heat map showing pairwise amino acid differences between HA sequences was generated (B), with purple indicating greater similarity and white indicating greater divergence. Predicted three-dimensional structures of the SG/16 HA (C) and pLM-generated HAs: 536 (D), 547 (E), 585 (F), 814 (G), 848 (H), 1025 (I), 1035 (J), were generated using Swiss-model (swissmodel.expasy.org), visualized in PyMol. Amino acid differences relative to SG/16 are shown in red, whereas conserved residues are shown in grey.

Predicted structures of the seven candidate HA proteins were modeled in PyMOL and compared with the SG/16 HA structure. Most of the amino acid substitutions in the HA candidates were localized to the HA head region surrounding the receptor binding site at positions 137, 158, 160, and 161 in antigenic site A, sites 175 and 176 in site B, and site 241 in site D (**Fig. 3C–J**).

### Candidate HA designs confer antigen-specific and broadly reactive antibody responses

To evaluate the antigenicity and protective efficacy of the seven candidate HAs, each antigen was expressed as recombinant HA (rHA) protein, formulated with AddaVax adjuvant, and administered to immunologically naïve DBA/2J mice (n = 11 mice/group) in a three dose intramuscular vaccination regimen at three week intervals. Serum samples were collected after each immunization to assess antibody responses. Following the final vaccination, spleenocytes from a subset of mice (n = 3/group) were analyzed for cellular immune responses, and the remaining mice (n= 8/group) were challenged intranasally with mouse-adapted SW/13 A(H3N2) to evaluate protective efficacy (**Fig. 4A**).

**Fig. 4.**
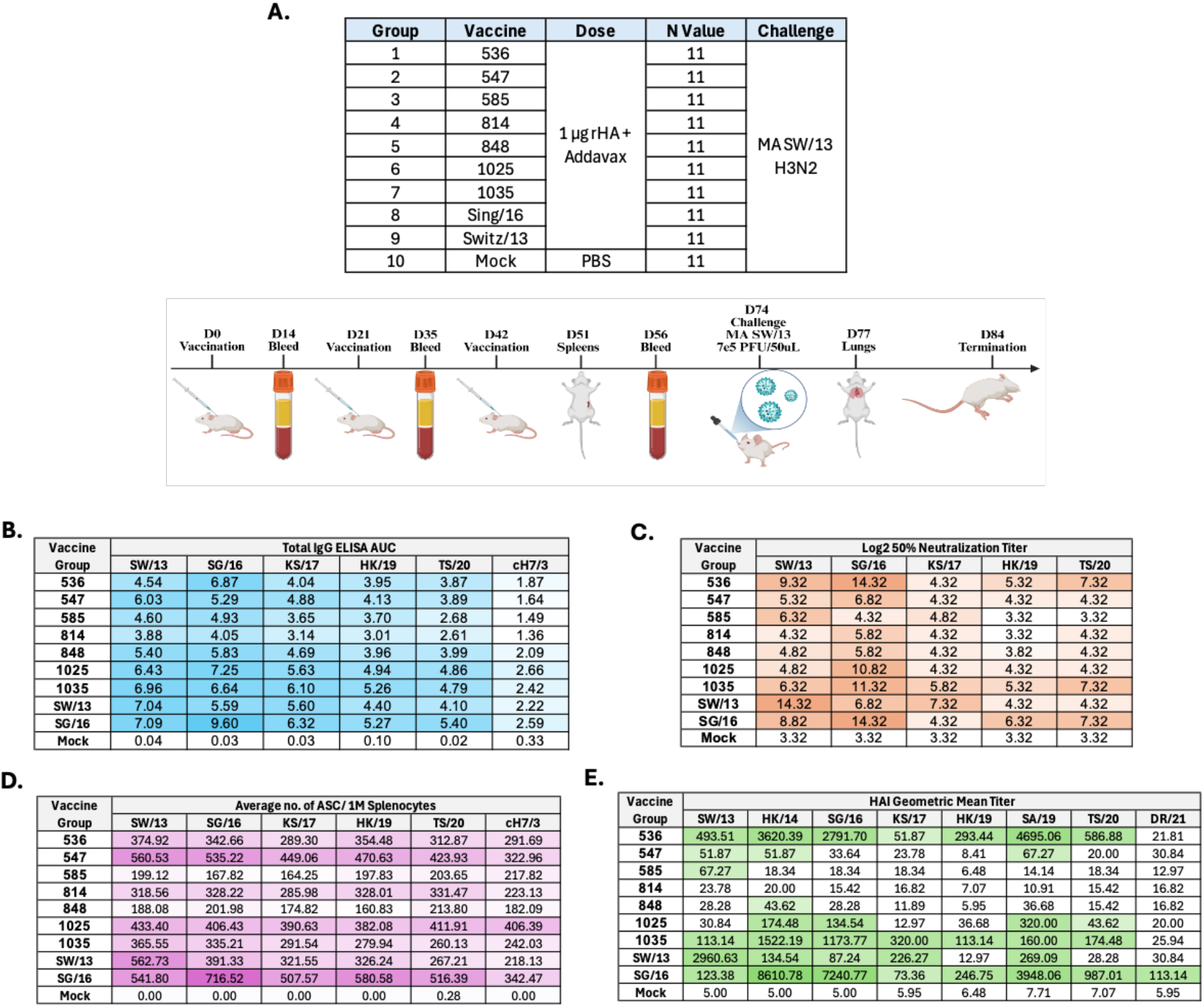
Mouse study outline and immunological evaluation of pLM-derived H3 HA vaccine candidates. Outline of the vaccination groups and study timeline (A). Heat maps depicting H3-specific IgG responses measured by ELISA area under the curve (AUC) using pooled sera collected after the third vaccination (B), log2 50% neutralization titers measured using pooled sera from each vaccine group (C), average frequencies of H3-specific antibody-secreting cells (ASCs) per 1 million input cells as determined by ELISpot from pooled splenocytes (n = 3/group) (D), and geometric mean HAI titers against historical A(H3N2) vaccine strains determined from individual serum samples (n = 8/group) (E). Increasing color intensity corresponds to higher IgG binding (blue), neutralization titers (orange), ASC frequencies (purple), and HAI titers (green).

Pooled sera collected after the third vaccination were evaluated for HA-specific IgG against rHAs representing World Health Organization (WHO) selected A(H3N2) vaccine strains circulating between 2013 and 2020 (**Fig. 4B**). The strongest responses were observed against SG/16 and the weakest against the chimeric cH7/3 rHA. Candidates, 1025 and 1035 elicited the highest HA-specific IgG responses, comparable to those induced by SG/16 rHA, while candidates 536, 547, and 848 generated intermediate responses similar to SW/13 rHA. Candidates 585 and 814 elicited the lowest overall IgG responses. Analysis of stalk-directed antibodies revealed a similar pattern, with candidates 848, 1025, and 1035 inducing the strongest anti-stalk IgG responses, whereas 536, 547, 585, and 814 elicited comparatively lower levels of stalk reactive antibodies (**Fig. 4B**).

Neutralizing antibody responses were assessed using pooled sera against a panel of A(H3N2) vaccine strains (**Fig. 4C**). Among the pLM-derived candidates, 536 and 1035 elicited the broadest neutralizing antibody titers across the panel, generating the highest titers against SG/16 and the future drifted HK/19 and TS/20 strains. Candidate 1025 produced strong neutralization against SG/16 but had reduced activity against the remaining viruses. Candidates 547, 585, 814, and 848 elicited lower neutralizing titers overall, although 50% neutralization titers remained 2- to 6-fold higher than those of mock-vaccinated animals. In contrast, the wild-type SW/13 elicited higher neutralization titers against SG/16 and the clade 3c.3a viruses SW/13 and KS/17 than the future drifted 3c.2a viruses HK/19 and TS/20. The SG/16 vaccine, however, generated broadly reactive neutralizing antibody responses across most strains in the panel, with the exception of KS/17 (**Fig. 4C**).

Cellular responses were assessed by quantifying HA-specific antibody-secreting cells (ASCs) using ELISpot assays (**Fig. 4D**). Candidates 547 and 1025 elicited the highest ASC frequencies across the H3 HA panel (∼400-600 ASCs per million splenocytes), comparable to those elicited by the SG/16 rHA, and also generated the strongest group 2 HA stalk-specific responses (∼300-400 ASCs per million splenocytes). Candidates 536, 814, and 1035 elicited intermediate ASCs frequencies (∼200-400 ASCs per million splenocytes), similar to the SW/13 rHA, whereas candidates 585 and 848 elicited the weakest responses, producing fewer than 200 H3-specific ASCs per million splenocytes across the panel (**Fig. 4D**).

Individual serum samples collected after the third vaccination were evaluated for hemagglutination inhibition (HAI) activity against a panel of A(H3N2) vaccine strains isolated between 2013 and 2021 (**Fig. 4E**). Among the seven candidates, 536 and 1035 elicited the broadest responses, generating mean sero-protective HAI titers against 7 of 8 strains. Candidate 1035 also elicted the highest HAI response against KS/17, while both candidates elicited significantly higher titers against HK/14 and SG/16 than candidate 814 (p < 0.05; **Fig. 4E**; **Supp. Fig. 1**). Candidates 547 and 1025 exhibited intermediate breadth, generating sero-protective titers against 3–4 strains, whereas candidates 585, 814, and 848 displayed limited breadth eliciting protective titers against at most one strain. In comparison, the wild-type SW/13 and SG/16 rHAs elicited sero-protective titers against 5 and 8 strains respectively. Notably, none of candidate HAs induced sero-protective HAI titers against the DR/21 isolate (**Fig. 4E**).

To investigate the reduced HAI reactivity against DR/21, ESM-2 sequence embeddings of the generated antigens were visualized alongside historical and contemporary A(H3N2) isolates using t-Distributed stochastic neighbor embedding (t-SNE) analysis^36^. The candidate antigens clustered with viruses circulating through approximately 2020, whereas later emerging isolates, including DR/21 were progressively separated in the embedding space. This separation was consistent with reduced HAI activity against DR/21, suggesting that the learned embeddings can provide a qualitative explanation for the temporal limits of sero-protection (**Fig. 5**).

**Fig. 5.**
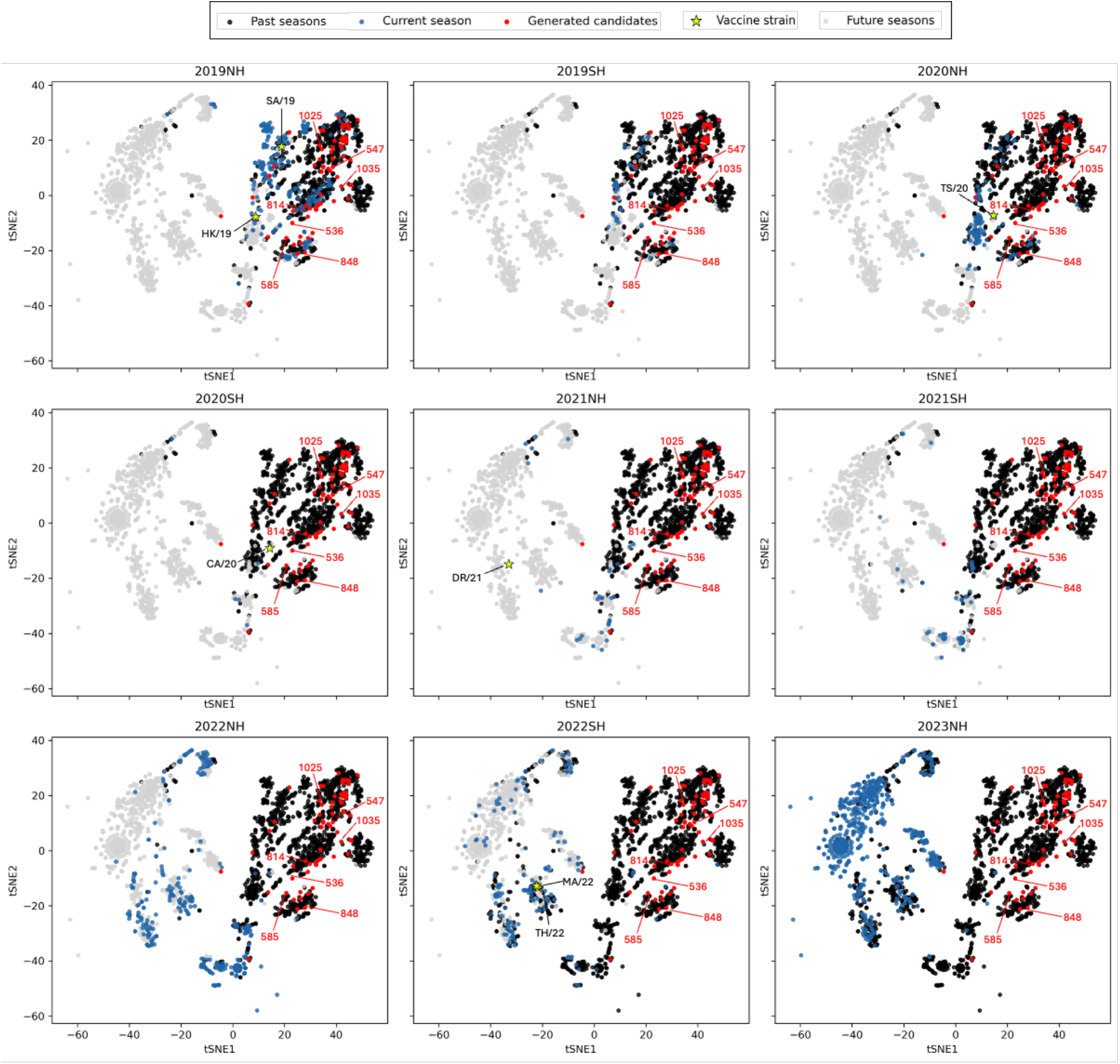
Selected candidates localize closely to isolates up to the 2020SH season in embedding space. t-SNE analysis of isolates embedded using HA1 fine-tuned ESM-2. Each plot emphasizes a different season (blue) in relation to past seasons (black), future seasons (gray), and generated candidates (red). The seven designs selected for experimental validation are marked, and vaccines strains from each season are also highlighted (star).

### Candidate HA designs protect mice against H3N2 challenge

Approximately three weeks after the third vaccination, mice were challenged intranasally with mouse-adapted SW/13 A(H3N2) virus, and monitored for weight loss, clinical disease, and survival (**Fig. 6**). Mice vaccinated with SW/13 rHA exhibited the least weight loss, averaging ∼3–5%, with minimal clinical signs. In contrast, mock vaccinated mice and those immunized with candidate 1025 experienced the most severe disease, characterized by ∼20% weight loss, elevated clinical scores, and ∼40-60% survival. Candidate 585 imparted partial protection with ∼80% survival following challenge. All remaining HA vaccine candidates limited clinical disease and conferretd complete protection, despite transient weight loss of ∼10-20% across days 3-4 of the infection. (**Fig. 6A–C**). Lung viral titers on day 3 post-infection demonstrated near-complete suppression of viral replication in mice vaccinated with candidates 536, 547, 585, 1035, SW/13, and SG/16 (**Fig. 6D)**. In contrast candidates 814, 848, and 1025 exhibited low, but measurable viral titers (∼10¹–10² PFU/g), whereas mock-vaccinated mice had the highest viral burden (∼10³ PFU/g).

**Fig. 6.**
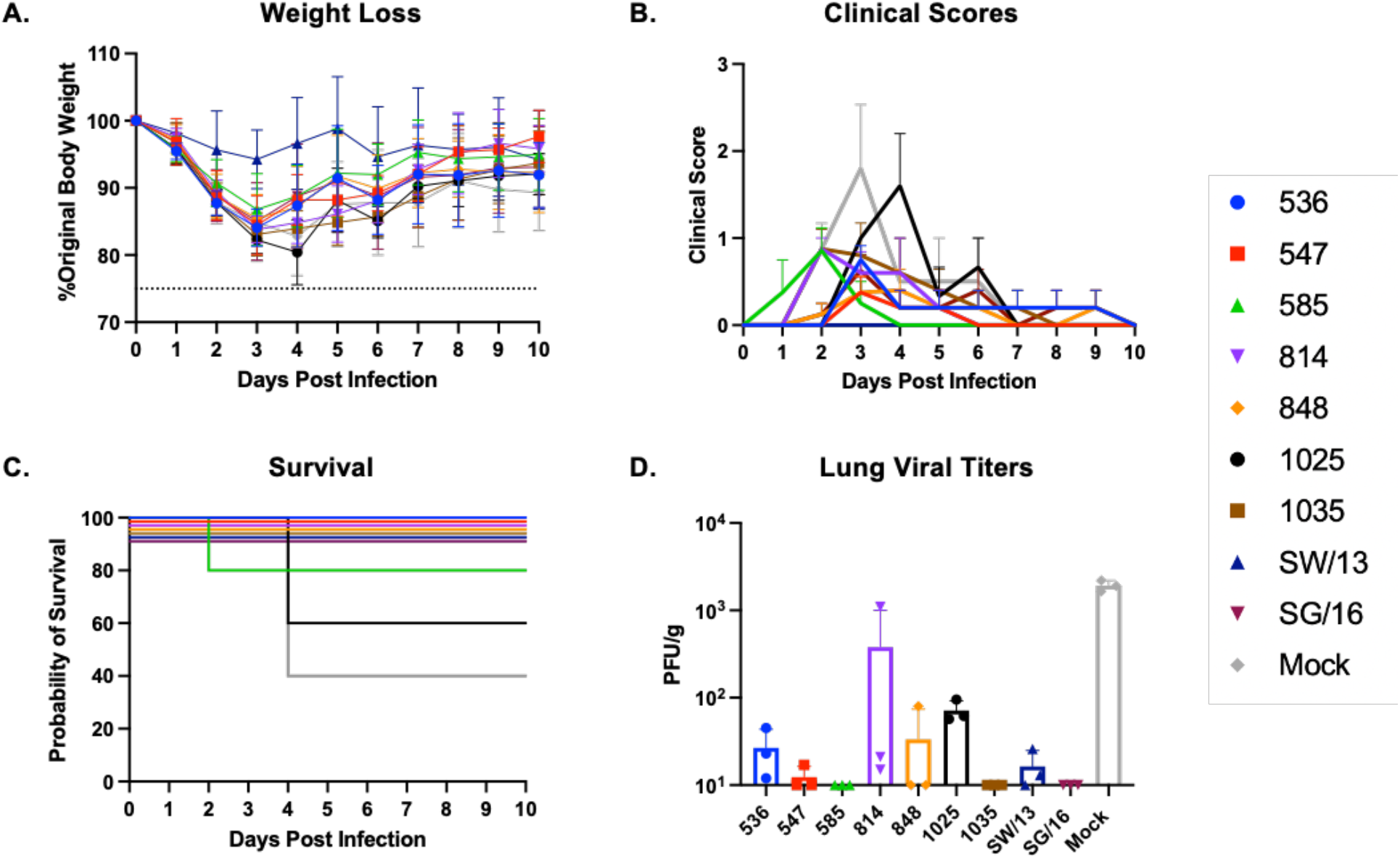
Mouse adapted SW/13 H3N2 influenza challenge. On day 74, the mice (n=8/group) were intranasally infected with MA SW/13 H3N2 virus, and monitored for weight loss (A), clinical scores associated with influenza infection (B), and survival (C). Three days post infection, lungs from a subset of mice (n=3/group) were collected and assessed for the presence of replicating virus (D). A Kruskal-Wallis ANOVA with Dunn’s multiple comparison test was used to determine statistical significance between each vaccine group (* = p<0.05, ** = p<0.01, *** = p<0.001, **** = p<0.0001).

## Discussion

Seasonal influenza vaccine effectiveness remains limited by the rapid antigenic evolution of circulating viruses, particularly within the A(H3N2) subtype. The findings presented here demonstrate that pLMs can generate HA antigens that elicit broad and protective immune responses against both contemporary and antigenically drifted A(H3N2) viruses. The integration of generative sequence design with antigenic prediction enabled the identification of vaccine candidates that preserved evolutionary plausibility while incorporating antigenic features associated with future viral lineages. Notably, sequence divergence from wild-type antigens was concentrated within antigenic sites surrounding the HA receptor-binding domain, suggesting that the model preserved key structural constraints while exploring sequence variation most likely to influence antigenicity. Together, these findings provide proof of concept that pLMs can generate biologically plausible antigens with broadened antigenic coverage beyond traditional surveillance-based strain selection.

Among the pLM-derived candidates, 536 and 1035 were the top performers, eliciting protective HAI responses, high frequencies of antigen-specific ASCs, and robust neutralizing antibody responses with cross-reactivity spanning contemporary 3c.2a and 3c.3a clades and future drifted strains. Sequence analysis revealed that both candidates contain features associated with multiple A(H3N2) lineages. A threonine substitution at position 128 in antigenic site A introduced a putative N-linked glycosylation motif characteristic of 3c.2a viruses^37^. Conversely, a lysine at position 160 in antigenic site B resulted in the loss of a glycosylation site associated with 3c.3a viruses^38^. Both candidates also have an asparagine at position 171, consistent with 3c.3a viruses and distinct from the lysine commonly found in 3c.2a strains, introducing a charge difference that may influence antigenicity^39^. Together, these sequence features likely contributed to the broad cross-clade immunity observed following vaccination.

Despite their breadth, the candidate HAs elicited limited sero-protective responses against the 2022 vaccine strain DR/21. Although the model captured antigenic variation present in TS/20, it failed to incorporate key amino acid substitutions associated with the 3c.2a1b.2a.2 lineage represented by DR/21, including residues 156S, 159N, 186D, and 190N within antigenic site B, and 53G in antigenic site C^40,41^. In particular, substitutions at positions 156 and 159 have been linked to major antigenic shifts in A(H3N2) viruses, and may have contributed to the reduced HAI activity against DR/21^42^. These findings suggest that the model’s design space did not fully capture emergent antigenic features associated with recently evolved A(H3N2) lineages, highlighting an opportunity for future refinement of the pLM-guided antigen design.

Several additional limitations should be considered. First, the antigenicity predictor was constrained by the availability of AHT-derived measurements, limiting evaluation on recent strain pairs. Second, because contemporary A(H3N2) viruses exihibit limited pathogenicity in mice without adaptation, protective efficacy could only be evaluated using the mouse-adapted SW/13 challenge virus^43^. Consequently, protection against future drifted viruses could not be directly assessed *in vivo*. Finally, pooled sera may have obscured inter-animal variabilility in immune responses, and our analyses focused primarily on antibody-mediated protection without defining epitope specificity or Fc-mediated effector functions.

Future studies should further refine the generative and predictive components of the framework. Incorporating structural information and broadly neutralizing antibody epitopes may expand antigenic coverage, while advances in pLM-based antigenicity prediction could reduce reliance on scarce HAI measurements through semi-supervised learning^26^. Experimentally, evaluating multivalent combinations of generated antigens and testing in more translationally relevant models, including ferrets, humanized mice, and animals with pre-existing influenza immunity, will be important for establishing clinical relevance.

Collectively, these findings establish pLM-guided antigen design as a promising strategy for developing broadly reactive influenza vaccines. Unlike surveillance-based approaches, pLMs leverage evolutionary and functional information encoded within large-scale sequence datasets to proactively design and prioritize novel antigens with broadened antigenic coverage. As increasingly powerful models become available, this framework could complement surveillance-based vaccine design and provide a generalizable approach for combating rapidly evolving pathogens.

## Methods

### Generative model development

#### Hemagglutinin sequence data curation

Influenza sequences can be sorted based on the representative influenza season in which they were collected: “Northern Hemisphere (NH)” (10/1/XX to 4/30/XY) and “Southern Hemisphere (SH)” (5/1/XY to 9/30/XY)^12^. Human influenza A(H3N2) HA sequences from the GISAID^28^ were collected from the start of 2013NH season (10/1/12) to the end of 2018SH season (9/30/18). The Biopython^44^ package was used to align the sequences with the full-length HA protein (566 amino acids) of A/Beijing/32/1992 (isolate ID: AAA87553) as a reference. MMseqs2^45^ was used to cluster the sequences at a 99% similarity threshold, in accordance with the WHO, which classifies influenza strains with more than four amino acid differences in the HA protein as epidemiologically significant^12^. Finally, the sequences were truncated to the HA1 region (amino acid sites 17-345), as this subunit contains the primary antigenic sites^46^. The final set of sequences were sorted based on the representative season in which they were collected.

#### Fine-tuning

ProGen2-medium^47^, a pretrained protein language model (pLM) with 764 million parameters, was selected as the foundation for adaptive fine-tuning on our HA1 sequence dataset. Fine-tuning was performed using HuggingFace *Trainer* for causal language modeling, with the Adam optimizer. A learning rate of 1×10^-5^ was used with the default linear learning rate scheduler in HuggingFace. A training batch size of 16 was used on NVIDIA GH200 machine. To prevent overfitting, the model is only trained for one epoch.

#### Evaluation

The data was split into training and validation sets using a time-series cross validation scheme in which the oldest season, 2013NH, is used to train the model, and validation is conducted on 2013SH. The validation set is then added to the training data, and the next season is used for validation. To assess model improvement on each fold, we calculated perplexity before and after finetuning. For a sequence *X* = (*x*_1_, *x*_2_, …, *x_n_*) of *n* tokens, the perplexity is calculated as follows:

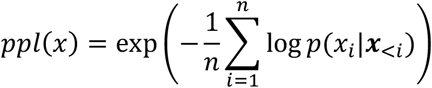

where *p*(*x_i_*|***x***_<*i*_) is the probability of the correct token *x_i_* conditioned on the preceding *i* − 1 tokens. Perplexity is a global metric used to quantify a language model’s understanding of its domain, and a lower perplexity indicates a better language model. To assess how the model understands “future” strains, the final model was trained on all seasons and evaluated on a holdout set of “future” sequences collected between 2019NH and 2023SH.

#### HA1 sequence generation and filtering

The final model was used to generate 1500 candidate HA1 sequences using a sweep over sampling temperature and nucleus sampling probability parameters: *T* ∈ { 0.8, 1.0, 1.2, 1.4, 1.6} and *P* ∈ {0.7, 0.9, 1.0}. Sampling temperature, *T*, effectively reshapes the model’s prediction confidence, with values below *T* = 1 sharpening the distribution toward the most confident amino acid predictions. Nucleus sampling, *P*, removes low-confidence amino acids from the predicted distribution, such that only the minimal set of amino acids contributing to *P* are selected from^47^. MMseqs2^45^ was used to cluster the designs at a 99% similarity threshold, and cluster representatives were dropped if they contained ambiguous residues or were shorter than 329 amino acids. A total of 356 candidate sequences remained after filtering. To quantify model uncertainty during sequence generation, the Shannon entropy was calculated at each sequence position. For an autoregressive language model, the predictive entropy at position *i* is given by

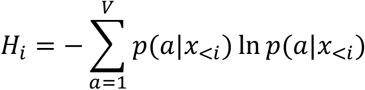

where *V* is the amino acid vocabulary size and *p*(*a*|*x*_<*i*_) is the probability assigned to amino acid *a* given the preceding sequence. Entropy is expressed in natural units (nats). The maximum entropy occurs when the model assigns equal probability to every amino acid, such that

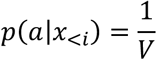

yielding

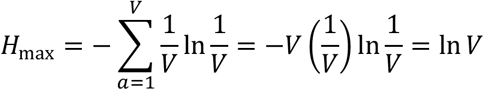

For a vocabulary size of *V* = 20 amino acids, the maximum predictive entropy is ln(20) = 2.996 nats. Predictive entropy was computed for every residue in every generated sequence and averaged across all generated sequences to obtain a positional entropy profile. To emphasize localized deviations in predictive uncertainty, the averaged entropy profile was detrended by subtracting a centered rolling median (window size = 11 residues), yielding a residual entropy profile used for visualization.

### Predictive model development

#### HAI dataset curation

The fine-tuning dataset was composed of hemagglutination inhibition (HAI) assay values collected from international reports and published articles that were compiled into a single benchmark dataset^29^. This dataset includes 3,698 A(H3N2) strain pairs collected between 1968 and 2010 with pre-processed antigenic distance calculations known as Archetti Horsfall Titers (AHT). AHTs use reciprocal HAI measurements between virus pairs, producing a symmetric metric that controls for non-antigenic effects such as viral red blood cell binding avidity^48,49^. This formulation has been shown to provide improved predictive performance for vaccine effectiveness relative to commonly used normalized hemagglutination inhibition titers (NHT)^49^. The antigenic distance (D_ab_) between strains *a* and *b* is defined as follows:

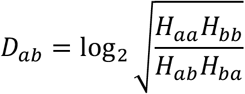

where the *H_ab_* HAI titer represents the maximum dilution of serum raised against strain *a* that is necessary to inhibit cell agglutination caused by strain *b*. Two viruses are defined as antigenic variants when the AHT is above 2; otherwise, they are considered antigenically similar. The dataset only denotes isolate years, so splits could not be denoted on a seasonal basis. Rather, the training set was composed from isolates collected prior to 2010, and a separate holdout set was created from all isolates collected in 2010. The model was trained solely using the data of past seasons to prevent data leakage that could inflate model performance.

To further evaluate the model’s generalizability, AHT measurements were computed from a dataset of 10,059 HAI measurements for isolates collected between 1968–2011^34^. After removing virus-antiserum pairs against which sequences were not found in the influenza genetic databases and removing pairs that match the training data, a total of 882 AHT measurements remained to test the model against.

#### Model architecture

ESM-2^32^, a 650M parameter model, was used as the backbone pLM. To improve model performance on the target domain^50^, ESM-2 was fine tuned under the masked language modeling objective (15% residue masking) using the previously curated dataset of HA1 sequences from 2013-2018. To prevent overfitting, a parameter-efficient fine-tuning strategy was implemented, Low-Rank Adaptation (LoRA)^33^ (rank = 32 and alpha = 64). The final model takes the HA1 sequences for both the virus and serum as input and independently passes them through the fine-tuned ESM-2 backbone. The sequence-level features for both sequences were extracted and the embeddings were subtracted as input to the downstream model to emphasize the encoded biological differences. The final model used a MLP constructed using the module *MLPRegressor* in Scikit-learn^36^. Hyperparameter tuning was performed using *GridSearchCV* with custom temporal splits as the cross-validation strategy, optimizing mean-squared error (MSE) across the entire validation scheme. The top model consists of two hidden layers with 128 and 64 neurons, respectively, using the ReLU activation function. Input features were standardized using a *StandardScaler*, and the network was trained with an initial learning rate of 1×10^-5^ and L2 regularization (alpha = 0.01). To ensure reproducibility, a fixed random state of 42 was maintained for each Python package across all simulations. For regression analysis, performance was evaluated using mean absolute error (MAE) and Pearson’s correlation coefficient (PCC) between predicted and observed AHT values. For classification analyses, AHT values were binarized using a threshold of 2 antigenic units (corresponding to a fourfold change in HAI titres). Virus–antiserum pairs with AHT > 2 were classified as antigenic variants (label = 1), whereas pairs with AHT ≤ 2 were classified as antigenically similar (label = 0)^29^. Model performance was then evaluated using accuracy, precision, sensitivity, specificity, F1 score, the area under the receiver operating characteristic curve (AUROC), and the area under the precision-recall curve (AUPRC).

#### Alternate models

The performance of the MLP model for A(H3N2) antigenic characterization was compared against alternative methods. Specifically, four ML methods were explored using available packages in Scikit-learn^36^: *KNeighborsRegressor*, *RandomForestRegressor, SVR,* and *GradientBoostingRegressor*. As before, the sequence-level embedding was extracted for both virus and serum sequences and the difference between the two was taken as the final embedding. For a fair performance comparison against the MLP model, hyperparameters of these models were optimized to minimize MSE using the same grid search scheme as previously described and performance was evaluated on the test set and the additional independent set.

#### Vaccine candidate selection

The final model was used for selection of candidate designs. Each design (n=356) was paired with each of seven selected “future” strains of interest: A/South Australia/34/2019 (SA/19), A/Hong Kong/45/2019 (HK/19), A/Tasmania/503/2020 (TS/20), A/Cambodia/e0826360/2020 (CA/20), A/Darwin/9/2021 (DR/21), A/Thailand/8/2022 (TH/22), and A/Massachusetts/18/2022 (MA/22). A selected influenza A(H1N1) strain, A/Victoria/2570/2019 (H1/VIC/19), served as a negative control group. The designs and test strains were passed through the model, and an AHT for each pairing was predicted. Designs that were predicted to be effective against all test strains (n=75) were down selected to maximize diversity within the primary antigenic regions. The important residue ranges were extracted and concatenated to generate antigenic-focused subsequences. Pairwise sequence dissimilarity was quantified using Levenshtein distance, yielding a full distance matrix. A greedy max–min selection strategy was applied to identify a subset of sequences with maximal diversity. The procedure was initialized by selecting the sequence with the greatest summed distance to all others. Subsequent sequences were iteratively chosen to maximize their minimum distance to the already selected set, ensuring broad coverage of sequence space. Selection continued until the desired number of sequences was obtained (n=30). The 30 H3 HA sequences were then aligned to H3 HA sequences from historical H3N2 vaccine strains that circulated from 2013 through 2021, and previously described COBRA H3 HA antigens J4, NG2, NG5, NG7, NG8^14,35^. Based on their phylogenetic relationships to these strains, 7 final HA sequences were down-selected for *in-vivo* characterization.

### t-SNE analysis

t-SNE visualization was used to qualitatively examine relationships among learned sequence representations, which were extracted from the ESM-2 model fine-tuned on influenza A(H3N2) HA1 sequences. Mean-pooled embeddings were generated for the top 70 HA designs scored by the predictive model together with historical and contemporary A(H3N2) HA1 sequences collected between 2013 and 2023. For visualization, the 1,280-dimensional embeddings were projected into two dimensions using the *TSNE* module in Scikit-learn^36^ with the cosine distance metric and random state set to 42.

### Experimental validation

#### Recombinant HA Synthesis

Full length HA sequences (566 amino acids) were generated by combining the HA1 region of each of the seven candidate HAs (amino acids 17-345) with the leader sequence (amino acids 1-16) and HA2 sequence (amino acids 346-566) derived from SG/16. Each sequence was then modified for recombinant protein expression by removing the HA transmembrane domain, and adding a T4 fold-on domain, an AviTag, and a 6x Histidine tag. The resulting constructs were cloned into a pcDNA3.1^+^ plasmid vector (Azenta Life Sciences, Burlington, MA). Each plasmid was transfected into HEK293T suspension cultures (Thermo Fisher, Waltham, MA) as previously described^51^. Supernatants containing soluble recombinant HA (rHA) proteins were harvested and purified via immobilized metal affinity chromatography (IMAC) using the C-terminal His tag. Protein concentrations were determined using conventional bicinchoninic acid (BCA) assays (Thermo Fisher). Protein purity and size were assessed by sodium dodecyl sulfate-polyacrylamide gel electrophoresis (SDS-PAGE), with 500 ng of each rHA loaded and electrophoresed at 200 V for 30 minutes, followed by Coomassie blue staining (Thermo Fisher). Each rHA was evaluated alongside 500 ng of trimeric His-tagged A/Texas/50/2012 control rHA (Sino Biological, Cat#40354-V04B)^35^. Observed molecular weights ranged from ∼60 to 90kd consistent with His-tagged trimeric HA. HA specificity was further confirmed by Western Blot analysis using a group 2-specific mouse monoclonal antibody (Immune Technology Corp, New York, NY; clone 34C9, Cat#IT-003-0423M13) diluted 1:2000, followed by incubation with a horseradish peroxidase (HRP)-conjugated goat-anti-mouse IgG secondary antibody (Southern Biotech, Birmingham, AL, USA) antibody diluted 1:4000. Signal detection was performed using Clarity ECL substrate (Bio-Rad, Hercules, CA, USA) and visualized on a Chemi-Doc imaging system (Bio-Rad) as previously described^35^.

#### Mouse Influenza Vaccination and Challenge

Influenza naïve female DBA/2J mice (n = 110), *Mus musculus*, 6-8 weeks of age, were purchased from The Jackson Laboratory (Bar Harbor, ME, USA). The mice were housed in microisolator caging, provided access to food and water ad libitum, and cared for following the USDA guidelines for laboratory animals. All procedures in this study were approved by the Cleveland Clinic IACUC (no. 2935). On day 0, 21, and 42 the mice (n = 11/group) were vaccinated with 1 μg of candidate recombinant HA (rHA) vaccine or wild-type rHA formulated with AddaVax adjuvant (InvivoGen, San Diego, CA) in a 50 μL volume of 0.9 % sterile saline. On days 14, 35, and 56 after the initial immunization blood was collected from each mouse. On day 51, spleens were collected from 3 mice per group. On day 74, the mice were intranasally challenged with 7×10^5^ PFU/50 μL of SW/13 A(H3N2) influenza virus. Following challenge, the mice were monitored for 10 days following infection for weight loss and clinical symptoms. On day 77 lungs were from 3 mice in each group, and on day 84 the remaining mice were humanely euthanized (n = 5 per group).

#### Viruses

Influenza A(H3N2) seed viruses were obtained from the Biodefense and Emerging Infections Research Resource Repository (BEI Resources), International Reagent Resources (IRR), or the U.S. Centers for Disease Control and Prevention (CDC), and were either propagated onsite or provided by Virapur. All viruses were passaged once in specific pathogen-free embryonated chicken eggs (AVS Bio, Norwich, CT) or Madin–Darby Canine Kidney (MDCK) SIAT cells. Following infection, virus-containing supernatants were collected and clarified by centrifugation to remove cellular debris, in accordance with WHO protocols. Hemagglutination titers ≥1:16, measured using 0.8% guinea pig red blood cells (Lampire Biological Laboratories, Pipersville, PA) supplemented with 20 nM oseltamivir carboxylate (Aobious, Gloucester, MA), were considered indicative of successful viral growth. After harvest, all virus stocks were aliquoted and stored at −80 °C for single-use applications.

The A(H3N2) viral panel used to assess the antigenic breadth elicited by the vaccines included the following historical vaccine viruses: A/Switzerland/9715293/2013 (SZ/13) egg passage 1 (EP1), A/Hong Kong/4801/2014 (HK/14) EP1, A/Singapore/IFNIMH16-0019/2016 (SG/16) EP1, A/Kansas/14/2017 (KS/17) EP1, A/South Australia/34/ 2019 (SA/19) EP1, A/Hong Kong/2671/2019 (HK/19) EP2, A/Tasmania/503/2020 (TS/20) EP2, A/Darwin/9/2021 (DR/21) EP1. Additionally, a mouse adapted A(H3N2) virus, A/Switzerland/9715293/2013 (SZ/13) EP2, was used to infect the mice on day 74 of the study.

#### Enzyme Linked Immunosorbent Assay (ELISA)

Immulon 4HBX 96-well microtiter plates (Cat. No. 3855, Thermo Fisher) were coated with recombinant hemagglutinin (rHA) proteins (SW/13, SI/16, KA/17, HK/19, TA/20, or cH7/3) at a concentration of 1 μg/mL in carbonate coating buffer (2.65 g Na₂CO₃, 2.1 g NaHCO₃, 450 mL diH₂O, pH 9.4). The cH7/3 is a chimeric HA with the H7 head of A/Anhui/1/2013 and the H3 stalk from A/Texas/50/2012^52^. A total of 100 μL of coating solution was added to each well, and the plates were incubated overnight at 4 °C in a humidified chamber. Following incubation, the plates were decanted, and 200 μL per well of blocking buffer (PBS containing 4% fetal bovine serum (FBS) and 0.05% Tween-20) was added to each well and the plates were incubated at 37 °C for 90 min in a humidified chamber. After blocking, the buffer was removed, and 100 μL of pooled serum samples from each vaccine group, serially diluted 3-fold in blocking buffer starting at 1:250, were added to each well and the plates were incubated at 37 °C for 90 min in a humidified chamber. After incubation, the plates were washed five times with wash buffer (PBS + 0.05% Tween-20). Subsequently, 100 μL of secondary antibody (goat anti-mouse IgG–HRP, 1 mg/mL; Cat. No. 1030-05, Southern Biotech), diluted 1:4000 in blocking buffer was added to each well, and the plates were incubated at 37 °C for 90 min in a humidified chamber. Following incubation, the plates were washed five times with wash buffer and 100 μL of substrate solution (1 mg/mL ABTS diammonium salt (Sigma), McIlvaine buffer (7.31 g Na₂HPO₄, 4.66 g C₆H₈O₇, 500 mL diH₂O, pH 5), and 3% H₂O₂) was added to each well and allowed to develop for 12–15 min at 37 °C in a humidified chamber. The reaction was stopped by adding 50 μL of 1% sodium dodecyl sulfate (SDS, Thermo Fisher) to each well. Absorbance was measured at 414 nm using a BioTek Epoch 2 plate reader (Agilent, Santa Clara, CA) equipped with Gen5 software. Optical density values were analyzed using GraphPad Prism Software (GraphPad, San Diego, CA). The area under the curve (AUC) was calculated for each vaccine group, and endpoint titers were defined as the reciprocal of the highest serum dilution yielding a signal greater than three times the mean of negative control wells.

#### Hemagglutination Inhibition Assay (HAI)

HAIs involving A(H3N2) isolates were conducted with a solution of guinea pig red blood cells (Lampire Biological Laboratories) diluted in PBS to 0.8%^14^. Prior to the assay, serum samples were treated with receptor-destroying enzyme (RDE) (Denka Seiken, Tokyo, Japan) according to the manufacturer’s instructions. In brief, RDE powder was resuspended in PBS and 300 µL was added to 100 µL of sera samples individualized in a 96-deepwell block. The mixture was then incubated overnight at 37 °C. After incubation, each sample is than heat treated in a 56 °C water bath for 45 min. The samples are then removed from the water bath and allowed to cool, once at room temp, 600 µL of PBS was added to each individual sample. After treating the sera samples with RDE, 50 µL of each sample was added to column 1 of a 96-well V-Bottom plate (Thermo Fisher), 25 µL of PBS was added to columns 2 to 11, and 50 µL of PBS was added to column 12. The sera samples were then serially diluted, 2-fold, across the plate by taking 25 µL of sera from column 1 and transferring it between each column across the plate until reaching column 11. Next 25 µL of influenza virus, diluted to 8 hemagglutinating units per 50 µL in PBS, was added to columns 1 to 11 of the plate. Plates containing A(H3N2) viruses were supplemented with 20 nM Oseltamivir Carboxylate (Aobious) and were allowed to incubate for 30 min at room temperature. After the incubation, 50 µL of 0.8% guinea pig RBCs were added to each well. The plates were incubated at room temperature for 1 h before being tilted to visually confirm the presence/absence of hemagglutination in each well. The HAI titer of the antibodies in the serum was then determined as the reciprocal of the dilution in the last well that had non-agglutinated RBCs. For this study, a “seroprotective” HAI antibody titer was defined as a titer ≥1:40^53^.

#### Microneutralization assay

A neutralization assay was used to determine the efficacy of vaccine induced antibodies to prevent virus infections in vitro. Neutralization assays were performed on pooled serum samples from each group of vaccinated mice, and the pooled samples were RDE treated prior to the assay. Each well of a 96-well flat bottom microtiter plate (Thermo Fisher) was loaded with 50µL of virus diluent (Dulbecco’s Modified Eagle Medium (DMEM)), 13.5% bovine serum albumin, 1% penicillin/streptomycin (P/S), 2.5% 1M HEPES buffer, and 2µg/mL TPCK trypsin (Thermo Fisher). The first column of each plate was supplemented with 40µL of virus diluent and 10µL of the pooled RDE treated serum. The serum was then serially diluted, 2-fold, by transferring 50µL from well to well across the plate, ending at column 10. Next, 50µL of virus that previously tittered to 100× of the 50% tissue culture infectious dose (TCID_50_) was added to each well of the plate, except column 12. Column 12 was instead supplemented with 50µL of virus diluent, so the wells of this column could serve as negative control wells. The plates were then incubated at 37 °C for 60 min. Following incubation, 100µL of MDCK-SIAT-1 cells diluted in virus diluent were added to each well at a concentration between 1.5 × 10^5^ to 2.0 × 10^5^ cells/mL, and the plates were incubated at 37 °C, 5% CO2 for 18 to 20 hr. After incubation, the plates were decanted, and each well was washed with 200µL of PBS. The PBS was then removed, and each well was fixed with a 4 °C solution of 80% acetone (Thermo Fisher) diluted in PBS. After 10 min, the fixative was removed, and the plates were allowed to dry for 20 mins at room temperature. The plates were then washed 3× with 200µL of wash buffer (PBS+0.03% Tween 20; Thermo Fisher) and 100µL of rabbit anti-influenza-A-NP (1 mg/mL; Cat. no. 40208-R061; Sino Biological) diluted 1:10,000 in blocking buffer (PBS+0.03% Tween 20 + 14.5% bovine serum albumin) was added to each well, and the plates incubated at room temperature for 60 min in a humidified chamber. After incubation, the plates were washed 3× with 200µL of wash buffer, and 100µL of goat anti-rabbit IgG HRP (horseradish peroxidase) (1 mg/mL; Cat. no. SSA003; Sino Biological) diluted 1:20,000 in blocking buffer was added to each well, and the plates incubated at room temperature for 60 min in a humidified chamber. The plates were washed 5× with 200µL of wash buffer, and 100µL of substrate (phosphate citrate buffer (Sigma), OPD tablets (Sigma), 0.05%, 30% H2O2) was added to each well. The substrate was incubated for 10-20 mins at room temperature, and once color change was detected in the positive control wells of column 11, 100µL of stop solution (0.5N H2SO4; Thermo Fisher) was added to every well. The plates were read at 490 nm on a BioTek Epoch 2 plate reader (Agilent) using Gen5 software. The 50% neutralization titer was determined by subtracting the average optical density (OD) values of the background signal, column 12, from the average OD of the virus control wells, column 11, and using that value as the 100% infection titer. The average background was also subtracted from the experimental wells, and the remaining values were compared with the 100% infection titers of the virus control wells. The 50% neutralization titer was then calculated for each group using a linear interpolation of the log2 serum titers above and below the 50% neutralization OD value. All assays were performed in duplicate, and the mean of the neutralization titers were calculated between the plates.

#### Antibody Secreting Cell Enzyme Linked ImmunoSpot (ELISpot) Assay

Spleens harvested after the third vaccination on day 51 were processed into single cell suspensions and examined for antigen-specific antibody secreting cells via the ELISpot assay^54^. In brief, In brief, pooled splenocytes from each group were stimulated using mouse-B-poly-S for 84-96 hours. Separately, 24 hours before the assay, MultiScreen 96-well clear plates with PVDF membrane filters (Cat. No. MSIPS45, Millipore Sigma) were coated with 1.5 μg/well of SW/13, SI/16, KA/17, HK/19, Tas/20, or cH7/3 rHA protein resuspended in PBS. Following stimulation, ∼300,000 pooled splenocytes from each group were resuspended in B cell medium (BCM)^15^, loaded into the first row of a deep well block, and serially diluted 2-fold to a final concentration of a 37,500 cells/100 μL. Next, the multiscreen plates were washed three times with sterile PBS, loaded with 100 μL of BCM/well, and 100 μL of each dilution of cells was added to the plates in triplicate. The plates were then incubated at 37 °C, 5% CO_2_ for 16-18h. Following incubation, the plates were washed three times (100 μL PBS supplemented with 0.1% TritonX; Thermo Fisher) and 5 times with PBS. Next, 50 μL of Goat Anti-Mouse IgG, Human ads-AP antibody, diluted 1:4000 in PBS+0.5% FBS (Fetal Bovine Serum) (Cat. No. 1030-04, Southern Biotech) was added to each well and allowed to incubate for 2 h at 37 °C. The plates were then washed and developed by adding 50 μL NBT/BCIP Substrate Solution (Nitro-Blue Tetrazolium and 5-bromo-4-chloro-3’-indolyphosphate) (Cat. No. 34042, Thermo Fisher) to each well. The substrate was allowed to develop for 10-15 min at 37 °C until spots formed in each well. Spots were imaged with a CTL ImmunoSpot^®^ analyzer (CTL) and counted using ImmunoSpot^®^ software version 7.0.38.2 Professional SC.

#### Influenza plaque assay

Plaque assays were performed using mouse lung tissue samples that were collected on day 77 to determine the amount of live virus present in the lungs of vaccinated animals following infection. In brief, 1 × 10^6^ MDCK epithelial cells (Sigma) resuspended in culture medium (DMEM, 10% fetal bovine serum, 1% P/S, 12.5 mL HEPES buffer, 12.5 mL 7.5% bovine serum albumin fraction V (Thermo Fisher)), were added to a 6-well culture plate (Thermo Fisher) and incubated at 37 °C for 18 to 22 h. Once the MDCK cells reached 90% to 95% confluency, the monolayers were washed 2× with DMEM supplemented with 1% P/S (DMEM + P/S) (Thermo Fisher). The media was removed from the wells, and 100 µL of each mouse lung tissue sample that had been homogenized, 70 uM filtered, then serially diluted 10-fold in DMEM + P/S, was added to each well of the plate. Each dilution was run in duplicate on each plate. The tissue samples were allowed to incubate at room temperature for 60 min, and the plates were agitated every 15-20 min to prevent drying. After incubation, the media the wells were washed 2× with DMEM + P/S. After washing, 2 mL of a mixture (1:1) of 2× Minimal Essential Medium (MEM), 1.6% agarose, 1 µg/mL of TPCK trypsin (Thermo Fisher) was added to every well. This mixture was allowed to solidify at room temperature, and then the plates were incubated at 37 °C 5% CO2 for 72 h. After incubation, the agarose gels were physically removed from each well, and the monolayers were fixed by adding 1.5 mL of 10% buffered formalin (Thermo Fisher) to every well. After 15 min, the fixative was removed and all the wells were stained using 1% crystal violet (Thermo Fisher) for 15 min. After staining, the plates were rinsed 5× with water and then allowed to air dry. The viral plaques on the dried plates were counted and recorded as the number of plaques present in the reciprocal of each dilution. The viral titers in the lung samples were reported as PFU/g of lung tissue for each sample.

## Data availability

The natural HA sequences and their metadata, including collection time and strain name, are from the GISAID^28^ and the IVR^55^ databases. The accession IDs for the sequences used in this study are available in our GitHub repository (https://github.com/IGlab-VUMC/FluGen and are organized under the respective models. The full, unfiltered dataset of HAI titer data used for training and evaluation of the antigenicity prediction models are available at figshare (https://doi.org/10.6084/m9.figshare.c.4961501)^29^. The additional, unfiltered holdout set can be found on Dryad (https://doi.org/10.5061/dryad.rc515)^34^. The original data presented in this study are made publicly available via the National Institutes of Health (NIH) ImmPort online database. The data will also be made available to qualified members of the scientific research community upon written request. Novel HA1 sequences generated in this study are available from the corresponding author upon reasonable request.

## Code availability

All code used for training and evaluating the pipeline will be made publicly available following journal acceptance and prior to publication at https://github.com/IGlab-VUMC/FluGen^56^. Model weights for the fine-tuned pLMs will also be available at https://huggingface.co/vrhoward/FluGen and https://huggingface.co/vrhoward/FluESM. Any additional data or code reported in this paper will be shared by the lead contact upon request.

## Ethics Statement

All research protocols, procedures, and humane endpoints involving mice were reviewed and approved by the Cleveland Clinic IACUC, protocol #2935 (most recent revision approved 06/13/2025). The Cleveland Clinic Florida Research and Innovation Center animal facility is a United States Department of Agriculture (USDA) inspected and American Association for the Accreditation of Laboratory Animal Care (AAALAC) approved research facility operating under Animal Welfare Assurance Number D21-01103. The facility is compliant with the standards for animal care outlined in the Guide for the Care and Use of Laboratory Animals as published in United States Department of Health and Human Services (DHHS) Publication Number (NIH) 85-23 (Eighth edition), and Public Health Service Policy on Humane Care and Use of Laboratory Animals, as Revised in 2015.

## Acknowledgements

We would like to thank the Influenza Reagent Resource (IRR), Influenza Division, WHO Collaborating Center for Surveillance, Epidemiology, and Control of Influenza, Centers for Disease Control and Prevention (Atlanta, GA, USA) for providing some of the A(H1N1) and A(H3N2) influenza viruses. We acknowledge all researchers at the originating and submitting laboratories that sequenced influenza viruses and made them available in GISAID and IVR databases. We would like to thank Spencer Pierce and Ron Nelson for producing, purifying, and quantifying the recombinant HA proteins. We also would like to thank Amanda Lynch and Jessica Mediana for their technical assistance carrying out the animal studies, and the Cleveland Clinic Animal Resource staff, technicians, and veterinarians for their excellent animal care. Some figures were created with BioRender.com

## Author Contributions

V.R.H., J.D.A., G.A.S., T.M.R., and I.S.G. conceptualized the study and experiments. M.T. conducted the animal work, collected samples, performed the serological and cellular assays, and wrote the methods sections on mouse vaccination, HAI, ELISA, ELISpot, and Neutralization assays. V.R.H and J.D.A. prepared the final figures, analyzed the data, and co-authored the manuscript with input from G.A.S., T.M.R., and I.S.G. All authors read and approved the final version of the manuscript.

