## Supplemental Figure 1 for "Design and characterization of broadly protective influenza A(H3N2) vaccine candidates using protein language models"

**Supplementary Figures.**

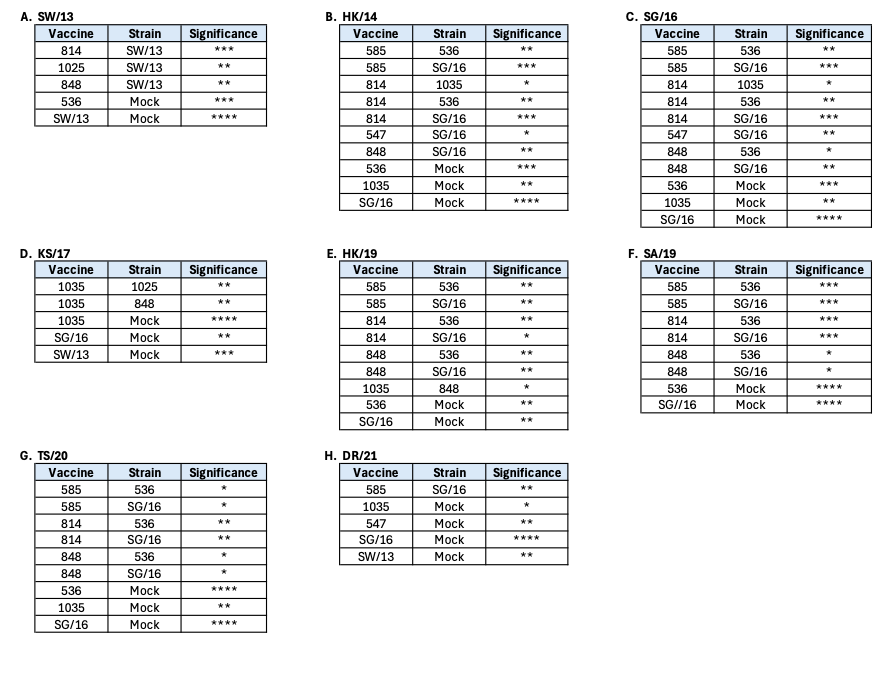

**Supp. Fig. 1.** Statistical significance of day 56 HAI titers. A Kruskal-Wallis ANOVA with Dunn’s multiple comparison test was used to determine statistical significance of the HAI titers between each vaccine group (* = p<0.05, ** = p<0.01, *** = p<0.001, **** = p<0.0001). Serum samples were tested against SW/13 (A), HK/14 (B), SG/16 (C), KS/17 (D), HK/19 (E), SA/19 (F), TS/20 (G), and DR/21 (H) H3N2 isolates.
